# Prioritising Autoimmunity Risk Variants for Functional Analyses by Fine-Mapping Mutations Under Natural Selection

**DOI:** 10.1101/2021.11.01.466789

**Authors:** Vasili Pankratov, Milyausha Yunusbaeva, Sergei Ryakhovsky, Maksym Zarodniuk, Estonian Biobank Research Team, Bayazit Yunusbayev

**Affiliations:** University of Tartu, Institute of Genomics, Tartu, 51010, Estonia; ITMO University, SCAMT Institute, Saint-Petersburg, 191002, Russia; Saint Petersburg State University, Institute of Translational Biomedicine, Saint Petersburg, 199034, Russia; University of Tartu, Institute of Bio- and Translational Medicine, Tartu, 50411, Estonia

## Abstract

Pathogens imposed selective pressure on humans and shaped genetic variation in immunity genes. This can also be true for a fraction of causal variants implicated in chronic inflammatory disorders. Hence, locating adaptive mutations among candidate variants for these disorders can be a promising way to prioritize and decipher their functional response to microbial stimuli and contribution to pathogenesis. This idea has been discussed for decades, but challenges in locating adaptive SNPs hindered its application in practice. Our study addresses this issue and shows that a fraction of candidate variants for inflammatory conditions evolved under moderate and weak selection regimes (sweeps), and such variants are mappable. Using a novel powerful local-tree-based methodology, we show that 204 out of 593 risk loci for 21 autoimmune disorders contain at least one candidate SNP with strong evidence of selection. More importantly, in 28% of cases, these candidates for causal variants colocalize with SNPs under natural selection that we fine-mapped in this study. Causal SNPs under selection represent promising targets for functional experiments. Such experiments will help decipher molecular events triggered by infectious agents, a likely early event in autoimmunity. Finally, we show that a large fraction (60%) of candidate variants are either hitchhikers or linked with the selected mutation. Our findings, thus, support both hitchhiking and natural selection models, with the latter having important practical implications in medicine.

## Introduction

Pathogens exerted strong selective pressure on human immune traits ^1,2^. Detecting the genetic footprints of these selection events can help us identify genotypes that were important for survival earlier in life and understand their later-life adverse consequences for immune-related diseases. This is a half a century old idea of antagonistic pleiotropy. It was initially proposed to explain ageing (Medawar 1952) but later was invoked to explain autoimmunity ^3,4^ and other human traits^5^. Indeed, genomic evidence suggests that immune-related genes were targets of natural selection ^6–10^. Accordingly, genetic risk loci for autoimmune diseases that harbour a subset of immune-related genes also bear signals of natural selection, represented by extended haplotypes ^10,11^. These findings, however, can be explained by two competing models: causal variants in these autoimmunity risk loci were driving selection signals, or they were hitchhiking with mutations undergoing selective sweep. If they were sweep-driving mutations, they are expected to have a strong effect on immunity traits. Such strong effect variants are ideal for detecting their function experimentally and, therefore, understanding their role in the pathogenesis of immune-mediated diseases ^11–14 4,15^. While the adaptive history of risk alleles for autoimmunity will offer new opportunities to understand their function under microbial invasion, it is unclear whether this model represents the prevailing scenario, i.e. the model that explains the observed overlap between autoimmunity risk loci and genomic signals of selection. Given that genomic signals of selection span up to hundreds of kilobases depending on the sweep intensity (strength of selection), there is ample room for causative SNPs to be hitchhikers ^3,14^. We start by outlining knowledge gaps in this area of research, highlighting the issues that hindered progress, and then describe our approach and major findings.

In humans, the role of pathogens as evolutionary agents of selection is poorly understood, and the evidence remains indirect, except for a few examples, such as malaria and cholera ^16^. This is in contrast to model organisms^17^ and plants^18^ that are suitable for experiments. The key work by Fumagalli et al. used a statistical approach to suggest that pathogens played an important role in shaping immune-related genes, including risk variants for autoimmune conditions ^1^. Similarly, another study indicated that Crohn’s disease risk loci overlap with the pathogen resistance signals in the human genome ^19^. More importantly, though, since mutations under selection and causal variants are hard to fine-map, there are only a few examples where we know that candidate causal variants had adaptive history and likely modulate resistance to pathogens. For example, Zhernakova et al. showed that candidate risk variants for Celiac Disease were under natural selection and that some of them correlate with a stronger response to bacterial ligands. This experimental evidence was taken to suggest that these disease risk variants likely confer resistance to pathogens ^20,21^. Another example is the genetic variant rs601338 in the FUT2 gene associated with Crohn’s disease ^22^. This variant was reported to confer a relative protective effect against norovirus infection depending on the pathogen genotype ^21–23^. It should be noted that these earlier works used haplotype-based signals of selection. In other words, haplotypes, which were constructed from chip genotyping data, provided only low SNP resolution, one SNP every 6-10 kb, on average. Low SNP resolution did not allow for teasing apart local genetic variation patterns, locating SNPs under selection, and comparing them with causal variants. Moreover, limited SNP resolution was further complicated by the lack of powerful methods to localize adaptive mutations ^24^, but there have been key methodological improvements recently ^25,26^. Another issue with earlier works is that they often used ad-hoc significance thresholds not suited to reject neutrality and did not control false signals due to population demography ^27^. Finally, earlier works detected only strong signals of selective sweeps in autoimmune risk loci ^10,11^. Such strong selective sweeps reduce genetic variation locally and represent a notorious obstacle for fine-mapping SNPs that drove selection. We refer to them as sweep-driving mutations throughout the text.

In contrast to strong selective sweeps, moderate sweeps, i.e. sweeps driven by mutations with weak and moderate selection coefficients, are harder to detect at the genome level. However, once weaker signals are identified, their sweep-driving SNP is easier to locate. Therefore, an important question is whether sweep-driving mutations related to immune phenotypes and inflammatory conditions evolved under strong, moderate, or weak selection regimens?

In this study, we report key progress in understanding the evolutionary history of the genetic component of inflammatory conditions. We used a recently proposed RELATE approach (Speidel et al., 2019) to build local trees for SNPs of interest and then the CLUES method to evaluate evidence for natural selection by computing the likelihood of selection scenario, coefficient of selection, and allele frequency trajectory ^28^. We applied this powerful approach to a large dataset of over 2400 fully sequenced genomes of Estonian Biobank donors. We show that whenever risk loci associated with inflammatory conditions show evidence for natural selection, selective sweep intensity was likely weak or moderate. This is an important finding since moderate and weak selection regimes (selection coefficients) imply that sweep-driving SNPs can be mapped. We then fine-mapped sweep-driving SNPs in 204 risk loci that showed evidence for selection. This allowed us to distinguish evolutionary scenarios underlying selection signals within analyzed risk loci, unlike previous haplotype-based approaches. Namely, evolutionary scenarios, where candidate causal SNPs likely evolved under positive or negative selection, were hitchhiking with adaptive mutation or were linked with an alternative haplotype to the selected one. Thus, the key contribution is that we identify SNPs that are promising candidates for experimental study of the causal SNP function under microbial exposure context. This is an important step to overcome current challenges with finding the disease variants and relevant physiological context to study their function in pathogenesis.

## Results

### Evidence for natural selection corrected for population demography

To test whether autoimmunity-associated loci evolved under natural selection, we started with 4331 SNPs that were reported by Farh et al., 2015 as potential causal variants for 593 unique GWAS hits (“index” SNPs) in 21 autoimmune conditions (Supplementary Figure 1). Additional 9151 SNPs were included from Estonian Biobank whole-genome sequences if they were in high LD (linkage disequilibrium measured as r2 ≥ 0.8) with at least one of the 4331 SNPs from Farh et al., 2015. This resulted in 13482 SNPs that we refer to as candidate causal SNPs throughout the text. To detect natural selection, we used the CLUES (Stern, Wilton, and Nielsen 2019) tool, which infers the allele frequency trajectory for a SNP of interest in modern DNA sequences. This is done by leveraging the properties of the local tree, which is a genealogy of sampled haplotypes at the tested SNP. Increasing and decreasing trajectories yield positive and negative selection scenarios. Local trees allow one to focus on a specific time interval when testing for selection and, besides, detect selection on standing variation. Moreover, CLUES not only computes the likelihood ratio of natural selection scenario over neutrality (logLR) but also infers the selection coefficient (s) (see Methods for the exact interpretation of s).

To choose a neutrality-rejecting threshold that takes into account the demographic history of the target population, we first assessed the distribution of logLR values using a simulation study. We simulated a demographic model representative of Estonian population history. The model parameters were based on the effective population size trajectory inferred from Estonian genome sequences using Relate (Speidel et al., 2019) (Supplementary Figure 2a, b). We then used the 95% percentile of the simulated distribution (logLR 1.59) as our neutrality rejection criteria (Supplementary Figure 2b).

We tested evidence for natural selection only for a subset of 9372 candidate SNPs (out of the 13482 candidate causal SNPs in high LD) if they mapped on RELATE-inferred local trees and passed all the filtering criteria for CLUES. These tested SNPs represent candidate causal variants for 550 risk loci. On average, each risk locus was represented by 17 tested SNPs (minimum one and maximum 158). Out of the 9372 tested SNPs, 1209 showed a median logLR value above the neutrality-rejection threshold of 1.59. These potential selection targets fall within 204 risk loci (on average ∼6 such SNPs per risk locus, minimum one and maximum 67). We, therefore, infer that ∼37% (204/550) of the risk loci for various inflammatory conditions contain at least one SNP that demonstrates evidence for natural selection.

### Strength of selection and mappability of mutations driving selection

Although traces of strong (high values of s) selective sweeps are easier to detect at the genomic level, fine-mapping sweep-driving mutations within such traces, i.e. sweep regions, is harder. Strong selection quickly brings the selected haplotype with linked SNPs to high frequency, thereby removing local SNP variation and leaving less time for recombination to restore flanking SNP variation that would be informative for fine-mapping. In contrast, weaker selective sweeps allow for more recombination and create local differences in SNP variation, making fine-mapping feasible. We, therefore, explored the distribution of selection coefficients (summarises strength of selection over a time period) among tested candidate SNPs (Figure 1a) to identify risk loci for which fine-mapping sweep-driving SNP is feasible. When all the candidate SNPs were considered, regardless of the significance of the selection signals, s ranged from 0.0013 to 0.013, which means that most of the observations correspond to weak and moderate selection sweep (Figure 1a), and only a few SNPs demonstrated strong selection sweep (Figure 1a). Here, we define strong, moderate, and weak selection events following the Schrider and Kern work ^29^. When we focused on SNPs with logLR≥1.59, we found that tested SNPs associated with the 204 risk loci mostly corresponded to sweep signals with moderate selection coefficients (orange circles at the central section, Figure 1a). These findings suggest that our candidate SNPs for downstream analyses changed in frequency relatively slowly, and there was time for recombination to counteract the sweeping effect. Our estimates, therefore, suggest that the local patterns of logLR variation within the reported 204 risk loci can be informative in prioritizing sweep-driving SNPs.

**Figure 1.**
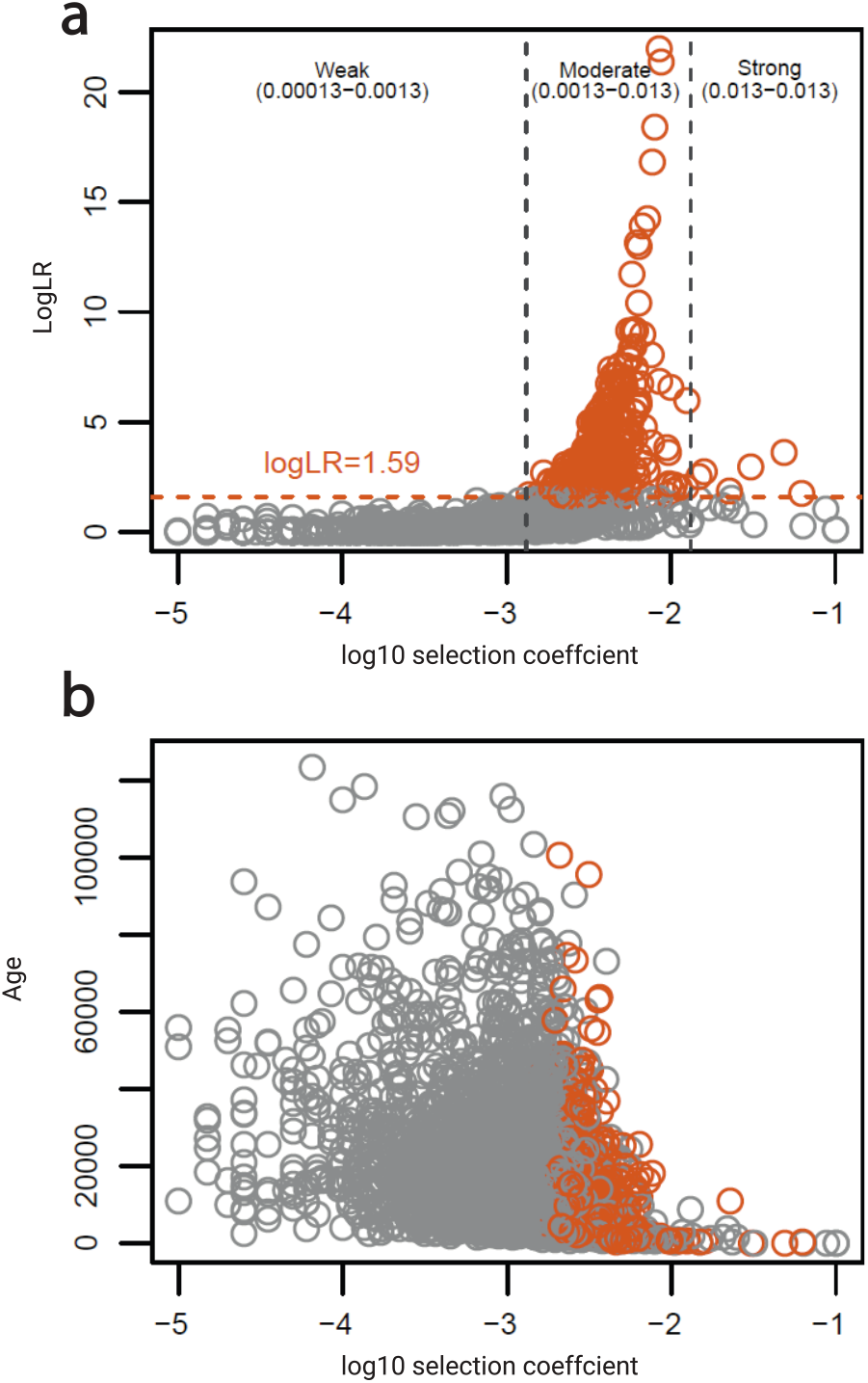
Joint distribution of logLR, selection coefficient, and allele age for 9372 candidate SNPs in 593 risk loci. Panel a graph shows the log-transformed likelihood ratio of the selection scenario versus the selection coefficient. The horizontal dashed line in orange indicates the neutrality rejection threshold. Vertical dashed lines (in grey) separate weak, moderate and strong selection coefficients, following the Schrider and Kern work ^29^. Corresponding untransformed selection coefficients are shown within parentheses. Panel b graph shows the distribution of derived allele ages for tested candidate SNPs and their selection coefficients. SNPs that have evidence for selection, i.e. logLR above neutrality rejection threshold, are shown in orange.

We note that fine-mapping selected SNP using local logLR variation attains very high resolution. This is because within a given haplotype block under selection, local trees can be inferred for very small chunks bounded by recombination. Consecutive local trees, inferred by Relate along such extended regions, would correlate in topology and branch-length, but they still slightly differ from one another due to historical recombination(s). Therefore, sweep-driving SNPs separated from linked neutral alleles by recombination are expected to yield higher logLR values. This principle was used in the original study ^28^ to fine-map the sweep-driving SNP.

Next, we inferred the approximate age of each tested SNP. We show that most of the candidate SNPs with evidence for natural selection (significant at logLR≥1.59) arose relatively recently in the history of human evolution, mainly after the out-of-Africa event (Figure 1b). It could be seen that there are only a few SNPs at logLR≥1.59 (on the bottom right) with a strong selection coefficient, and they all correspond to relatively recent allele ages. This agrees with the expectation that adaptive alleles with stronger selection coefficients would quickly reach fixation and be missed by sweep detection analysis.

### Prioritizing SNPs under selection and colocalizing selection targets with candidate causal SNPs for the autoimmune conditions

It was shown in the original study that the logLR could be used to fine map the sweep-driving SNP ^28^, and the accuracy is only limited by the extent of linkage disequilibrium (See Figure 8A in ^28^). We computed logLR values for a consecutive set of candidate SNPs bound by recombination within each risk loci with evidence for selection (logLR≥1.59). The highest logLR values were then taken to pinpoint the SNP(s) that likely drove the selection signal. We then asked whether these predicted sweep-driving SNPs matched the likely causal variants within the risk locus. For that, the highest logLR values were compared with the highest PICs (Probabilistic Identification of Causal SNPs) scores; the latter estimates the probability of the candidate SNP to be causal ^30^. We identified a range of scenarios in this way, and results for all the 204 risk loci are available in Supplementary data 1. For example, the most likely sweep-driving SNP and its positively selected allele matched the most likely causal variant and its effect allele (Figure 2a), or cases where the prioritized sweep-driving SNP had derived (selected) allele, but the effect allele for the top causal variant was ancestral (Figure 2c). These two scenarios identify SNPs that are promising targets for downstream functional analyses. However, we often found that the prioritized sweep-driving SNP was different from the top candidate causal variant, i.e., sitting nearby or far away from it (Figure 2b, d). In such cases, we compared the haplotype phase to learn whether it corresponded to genetic hitchhiking. Figure 2b demonstrates an example of genetic hitchhiking, where the disease allele (effect allele) for the top candidate SNP was on the same haplotype with the selected allele driving the sweep. Figure 2d exemplifies the alternative scenario, where the effect allele for the most likely causal SNP was on the different haplotype to the selected allele driving the sweep signal.

**Figure 2.**
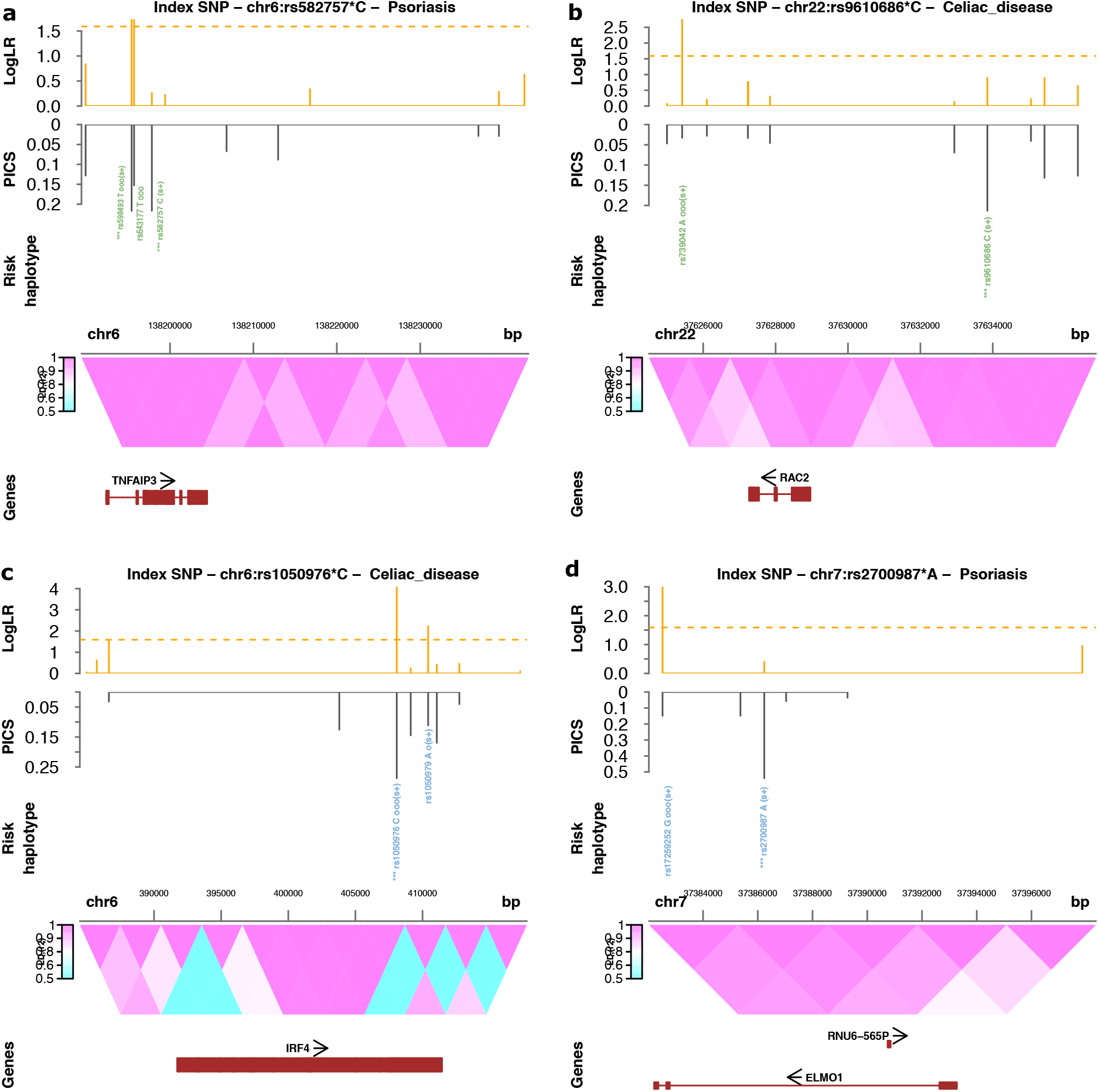
Examples of candidate risk SNPs that are hitchhiking and that match prioritized sweep-driving mutations. Panel a shows an example of risk loci, where one of the top candidate SNPs with the highest PICs score (^***^rs598493 T ooo(s+)) is likely a sweep-driving mutation. In panel b, top candidate SNP (^***^ rs9610686 C (s+)) is hitchhiking with nearby variant rs739042 A ooo(s+), the sweep-driving mutation. Five graphs (from top to down) represent five layers of annotation for candidate SNPs within the analysed risk loci, marked by index SNP. The first graph on the top shows logLR values. The horizontal dashed line in orange separates logLR values (above neutrality threshold), suggesting evidence for selection. The second graph depicts PICs values for each candidate SNP reported in Farh 2015. The third line from the top shows candidate SNP rsID along with the disease effect allele. The derived SNP allele is shown in green and in blue, when ancestral. Most likely candidate causal SNPs, which have the highest PICs values, are indicated with three asterisks (^***^). SNPs with evidence for selection (logLR 1.59) are indicated with “o” suffix, and “ooo” denotes the most likely sweep-driving mutation. s+ and s-indicate positive and negative selection coefficients, respectively. The fourth graph shows a heatmap of pairwise linkage disequilibrium between SNPs, measured in r^2^. The fifth graph on the bottom depicts gene annotations from the Ensembl genome database (version 87, Human genome build GRCh37).

Our high-resolution analysis suggests that some of the 204 risk loci contain multiple sweep signals that represent a composite of evolutionary scenarios. Since selection signals often appear as an extended region of depleted diversity, such complex scenarios could have been missed by previous haplotype-based tests. For example, a causal SNP might appear as the most likely adaptive SNP locally, but sequence data combined with a Relate/CLUES approach can distinguish stronger sweep signals nearby, attaining higher resolution (Supplementary Figure 3). In this case, one cannot easily rule out the hitchhiking scenario. We distinguish such complex scenarios from clear examples of hitchhiking and adaptive history for candidate causal SNPs. In Figure 3, we summarised all the possible scenarios, including complex ones and the number of times they were encountered (See **Supplementary Data 1** for the annotation graphs for the 204 risk loci used for this summary). Thus, our high-resolution analysis shows that genomic signals of natural selection in disease risk loci stem from various evolutionary scenarios that could not be recognized in earlier studies.

**Figure 3.**
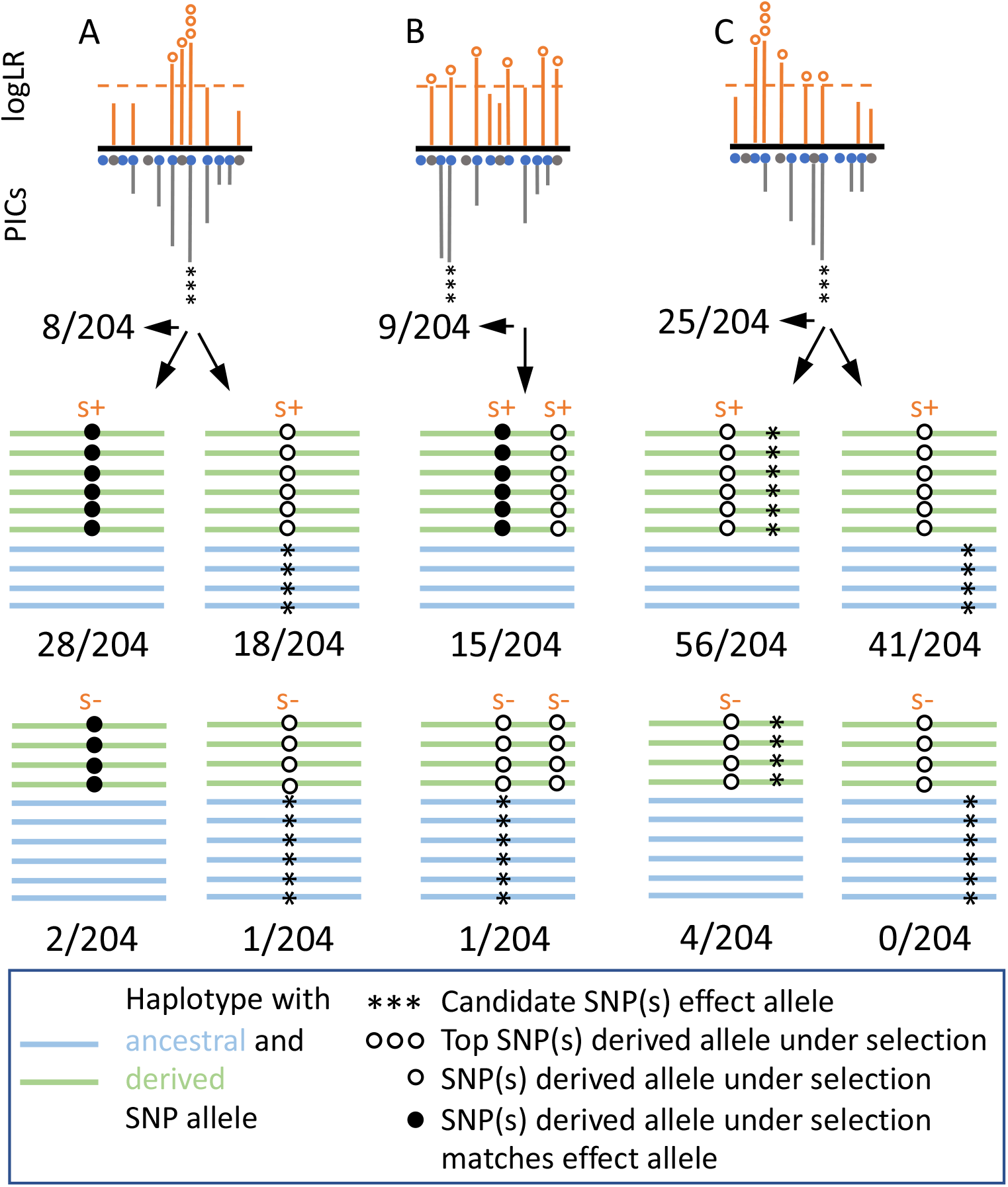
Evolutionary scenarios for candidate causal SNPs uncovered using logLR fine-mapping. The top line shows three basic scenarios (A, B, C) discernible for the 204 disease risk loci with a signal of selection: A) the top candidate causal variant (with the highest PICs value) matches the likely sweep-driving mutation (SNP with the highest logLR value); B) the top candidate causal variant shows evidence for natural selection (logLR 1.59), but might be hitchhiking since linked with a stronger target of selection; C) the top candidate causal variant is hitchhiking with nearby sweep-driving mutation. Note that the effect allele of the top causal variant might be derived and be identical with the sweep-driving allele. Alternatively, it can be ancestral and likely not experienced selection. We, therefore, further distinguish five sub-categories that are depicted on the second and third rows. The second and third rows show instances when the sweep-driving allele has the positive selection and negative selection coefficients. **Supplementary Data 1** contains full annotation graphs for the 204 risk loci classified in this figure.

### Revisiting previous studies and the role of genetic hitchhiking

It has been long hypothesized that causal alleles for inflammatory conditions were driven by genetic hitchhiking. With a few empirical examples, the role of this evolutionary scenario is poorly understood ^14^. We analyzed the largest collection of risk loci to date and found that genetic hitchhiking (56/204, Figure 3) is more frequent than adaptive history when there is a genomic signal of natural selection (28/204, see Figure 3). We, therefore, revisited earlier studies that reported adaptive history for disease alleles, but which had limited resolution in terms of data used (haplotype data) and methodology. For example, one of the pioneering works suggested that the Celiac Disease rs3184504*A variant in the SH2B3 gene had adaptive history and possibly played a role in pathogen resistance ^20^. Zhernakova et al., 2010 showed that this SNP variant demonstrated strong evidence for positive selection based on the haplotype-based iHS statistics (iHS=-2.597, p = 0.009). Moreover, donors with this risk allele demonstrated stronger, dose-dependent expression of 3 inflammatory cytokines in response to bacterial ligands in monocytes ^20^. We reanalyzed this SNP region and confirmed the strong evidence for positive selection in this SNP region (rs3184504, logLR=16.8512) (Supplementary Figure 4). However, in our sequence data, we found novel variants in strong LD with the rs3184504 SNP (r2≥0.9-1) that demonstrated comparable (rs7310615, logLR=16.8512) and even stronger evidence for positive selection (rs653178, logLR=21.9757) (Supplementary Figure 4). Thus, our analyses suggest that there is a better candidate for the true adaptive SNP in this risk loci than the rs653178 variant, and further work is needed to verify the true adaptive scenario for this genomic region.

We next revisited one of the most comprehensive studies that suggested natural selection history for genes implicated in inflammatory diseases ^11^. Raj et al. tested 416 risk SNPs for inflammatory conditions ^11^ using the haplotype-based iHS-score. They reported 21 SNPs with extreme iHS-scores (|iHS| ≥ 2) and suggested a positive selection scenario for them. We analyzed these SNP regions and found that for most of these loci, CLUES (14 out of 21 SNPs had logLR<1.59) yield surprisingly weak support for natural selection, with logLR varying between 0.04-1.44. Only 7 out of the 21 previously reported SNPs showed evidence for natural selection at logLR≥1.59 (Table 1).

**Table 1.** Revisiting published evidence on selection signals in inflammatory disease risk loci. ^a^ CLUES-based selection tests from this study. Evidence for selection, logLR 1.59 ^b^ Selection test based on iHS from Raj et al 2013. Evidence for selection, |iHS 2|

| SNPs reported<br>in Raj et al 2013 | Chromosome | Position<br>(GRCh37) | median logLR<sup>a</sup> | s | CLUES<br>tested<br>derived<br>allele | Signals of<br>selection<br>confirmed<br>with CLUES | iHS<sup>b</sup> |
| --- | --- | --- | --- | --- | --- | --- | --- |
| rs1167796 | chr7 | 75173180 | 0.4036 | 3E-04 | A | Not confirmed | 5.64 |
| rs3131379 | chr6 | 31721033 | 0.0429 | 8E-04 | A | Not confirmed | -3.18 |
| rs17696736 | chr12 | 1.12E+08 | 13.192 | 0.006 | G | Confirmed | -3.10 |
| rs281379 | chr19 | 49214274 | 1.4498 | NA | G | Not confirmed | 2.98 |
| rs3184504 | chr12 | 1.12E+08 | 16.851 | 0.008 | T | Confirmed | -2.97 |
| rs11962089 | chr6 | 1.06E+08 | 0.7165 | 0.004 | A | Not confirmed | 2.81 |
| rs6822844 | chr4 | 1.24E+08 | 1.2528 | 0.003 | T | Not confirmed | -2.82 |
| rs4308217 | chr3 | 1.22E+08 | 3.1177 | 0.007 | G | Confirmed | -2.73 |
| rs12638253 | chr3 | 1.57E+08 | 6.6191 | 0.01 | G | Confirmed | -2.71 |
| rs17810546 | chr3 | 1.6E+08 | 3.02 | 0.005 | G | Confirmed | -2.66 |
| rs1132200 | chr3 | 1.19E+08 | 0.3386 | 0.001 | G | Not confirmed | -2.68 |
| rs307896 | chr19 | 47661493 | 4.9529 | 0.004 | A | Confirmed | -2.62 |
| rs2188962 | chr5 | 1.32E+08 | 3.6903 | 0.004 | T | Confirmed | -2.60 |
| rs12708716 | chr16 | 11179873 | 0.2297 | 6E-04 | G | Not confirmed | 2.32 |
| rs11755393 | chr6 | 34824636 | 0.1667 | 0.001 | C | Not confirmed | -2.28 |
| rs10210302 | chr2 | 2.34E+08 | 1.0041 | NA | C | Not confirmed | 2.18 |
| rs415890 | chr6 | 1.67E+08 | 0.1423 | -4E-04 | T | Not confirmed | 2.15 |
| rs3129934 | chr6 | 32336187 | 0.0429 | 8E-04 | A | Not confirmed | -2.12 |
| rs10440635 | chr5 | 40490790 | 0.9178 | NA | G | Not confirmed | 2.03 |
| rs3745516 | chr19 | 50926742 | 0.1126 | NA | A | Not confirmed | 2.01 |
| rs2838519 | chr21 | 45615023 | 0.2896 | -8E-04 | A | Not confirmed | 2.00 |

The positive and negative selection that we inferred represents only two possible scenarios for the complex trait evolution. Both theory and experimental evolution suggest that genetic loci involved in resistance to pathogens might follow a frequency-dependent selection ^31^. This selection regime creates the so-called balancing haplotypes. We, therefore, asked if any of the risk loci covered by the sequenced 9372 SNPs overlap with balancing haplotypes in the human genome. We screened haplotypes with strong evidence (top Beta scores) for balancing selection in European populations ^32^ and found 565 candidate SNPs (in strong LD with eight index SNPs) that fell within the balancing haplotypes (Supplementary Table 2). These SNPs were missed by the logLR approach. Thus, combined with the 204 loci under positive and negative selection, eight loci under balancing selection suggest that around ∼35% of the risk loci (212 out of 593) for inflammatory conditions contain variants that were important for human adaptation.

## Discussion

Pinpointing causal mutations for complex diseases is challenging, and one underutilized principle is looking for SNPs under natural selection. The rationale is that such variants must strongly affect the trait to be picked by natural selection. Our work was motivated by this idea, and we addressed several issues that hindered its application. We used a novel methodology to show that approximately 35% (212 out of 593) of the risk loci for various inflammatory conditions contain mutations that were important for adaptation. This fraction is substantially larger than reported before (21/416 ∼ 5%) for inflammatory conditions ^11^. As discussed below, we attribute this to a better power of detecting weaker signals of selection with the Relate/CLUES approach.

We next asked if we could fine-map sweep-driving mutations in these sweep regions. We showed that mapping is possible by demonstrating that most candidate SNPs within disease risk loci corresponded to weak and moderate sweep scenarios (selection coefficients). Only a few SNPs demonstrated signals of strong selection (Figure 1a). Weak and moderate selective sweeps leave more time for recombination to create increasing nucleotide diversity flanking the sweep-driving mutations. This is in contrast to strong sweep signals that leave little information for fine-mapping and that were reported by earlier studies ^10,11^. The inferred weaker selection signals for candidate SNPs also explain why previous studies found fewer sweeps among risk loci ^10,11^. That said, we recognize that our study can miss even weaker signals of selection that correspond to highly polygenic sweeps for immune-related traits ^33^. We also note that our neutrality rejection approach helped us detect more selection signals at the cost of incurring small (5%) amounts of false rejections that can be explained by target population demography (Supplementary Figure 2a). This is a small cost if the goal is to have more targets for experimenting in the future.

The selective sweep scenarios suggested by our analyses motivated us to attempt to fine-map the sweep-driving SNP among the candidate SNPs. By inspecting local logLR variation for candidate SNPs, we show that in ∼28% of cases (57/204), candidate causal SNP matched with the likely sweep-driving SNP. Here, we note that the fine-mapping resolution will be limited by linkage disequilibrium in the focal loci like with any fine-mapping approach. Therefore, we prioritize more than one tightly linked SNP to be a sweep-driving mutation in some cases.

Interestingly, more candidate SNPs (∼27%, 56/204) were hitchhikers or were linked with the non-selected haplotype (∼20%, 41/204). Our study is the first comprehensive survey to provide empirical evidence on the hitchhiking scenario to the best of our knowledge. Hitchhiking has been long hypothesized for genes implicated in inflammatory conditions but there were only a few examples to support this model ^34^.

In contrast to more frequent hitchhiking, the natural selection scenario that we discovered for 57 candidate causal SNPs can help address some of the challenges in modern medical genomics. Namely, help in fine-mapping and finding the physiological context ^35^. Indeed, most causal variants for chronic inflammatory diseases hide in the non-coding DNA, hindering their fine-mapping within risk loci ^30,36^, especially when the relevant physiological context is unknown. As briefly mentioned before, the idea is that natural selection picked adaptive variants if they had a strong effect on the molecular traits serving better survival. In our case, we expect these molecular traits to be involved primarily in immune response or in immunometabolism, which ensures immunity at the organismal level ^37^. Hence, if adaptive mutations contribute to disease pathogenesis, the strong effect on the underlying molecular trait must be easier to detect and study experimentally. Therefore, when considering multiple candidates across risk loci or within risk loci, variants with adaptive history can be promising targets to prioritize for functional study. Our work, thus, represents an important milestone in the overall fine-mapping effort in the field.

More importantly, the adaptive model predicts that pathogens or structurally similar symbionts represent a relevant trigger to elicit the regulatory function of candidate causal SNPs for inflammatory conditions. The regulatory function of such SNPs in immune cells can often go undetected and stay “silent” ^38^ unless immune cells are triggered by the stimuli, which is specific for the disease pathogenesis^39^. Indeed, it has been highlighted many times that external stimuli and stimulus-specific regulatory responses are important for discovering immune-related genetic variants in inflammatory conditions ^40,41^. Our findings inspired by the adaptive hypothesis offer promising candidates to test the utility of the microbial exposure context in studying causal variant function in autoimmunity.

Microbial exposure has long been used to trigger an autoimmune response in animal disease models ^42^. There are numerous examples where pathogens (viruses, bacteria, and parasites) are known to trigger human autoimmune disease and exacerbate it further ^43^. For example, tonsillar infection with Streptococcus pyogenes has been known for decades to trigger and exacerbate psoriasis skin lesions^44–46^. Even in Celiac Disease, aberrant response to dietary gluten is initiated by a viral infection ^47^. Despite the accumulated evidence, infectious triggers of autoimmune diseases, such as, for example, Herpesvirus, are known to be common in a healthy population and do not lead to autoimmunity. Therefore, the major knowledge gap is whether pathogens interact differently with individuals having risk alleles to autoimmune diseases, and that results in autoimmunity? Our study offers an actionable list of 57 candidate risk SNPs to ask this question for various autoimmune diseases using experiments. Such experiments can shed light on molecular pathways that microbial exposure initiates in carriers of risk SNPs and illuminate early events in pathogenesis. The latter is a poorly understood aspect of autoimmunity with promises for treatment.

## Methods

### Genetic loci associated with autoimmune diseases

In this study, we retrieved data on 613 (593 unique loci if we remove duplicated between diseases) genomic risk loci associated with 21 autoimmune disorders, for which the previous study prioritized 4950 (4331 unique) potentially causal SNPs (“candidate SNPs”), each annotated with a PICs (Probabilistic Identification of Causal SNPs) score. PICs score reflects the probability of the candidate SNP to be causal given the haplotype structure and observed pattern of association at the locus ^30^ (Supplementary Table 1, Supplementary Fig 1). As we are interested in evidence for positive selection not only at the potential candidate SNPs but also at linked loci, in addition to the 4331 unique candidate SNPs from Farh et al. 2015 we selected 9151 SNPs in high LD (r2 ≥ 0.8) with at least one candidate SNP among the Estonian Biobank samples with whole-genome sequences available (see below for details on the dataset used). As a result, we started our analysis with a list of 13482 variants.

## Data availability

The sequencing data are available upon request. The application procedure to access the data can be found under the following link: https://www.geenivaramu.ee/en/biobank.ee/data-access.

### Evidence for natural selection based on local trees

A typical genomic risk locus for a complex disease usually spans several tens of kilobases and contains multiple potentially causal SNPs. Such genomic regions can be subdivided into smaller chunks bounded by recombination events that occurred during the evolutionary history of the risk loci. Each such chunk has a genealogy that correlates with nearby subregions (due to LD. Such genealogies or local trees reflect the full history of the genomic subregion and candidate SNPs within it (Supplementary Fig 1). If the candidate SNP had a selective advantage in the past, one would expect the selective sweep to distort the local tree and deviate from neutrality expectations in terms of its topology and branch lengths. The extent of distortion can be translated into the likelihood of natural selection and the competing neutrality scenarios. This is the most accurate way to test for evidence of selection. Both elements of this powerful concept are implemented in the recently proposed RELATE approach ^48^ to infer local trees and the CLUES algorithm to compute the log-likelihood of competing models ^28^. We used this powerful approach to test evidence for natural selection for candidate SNPs within the 593 risk loci.

Specifically, we first inferred local trees for subregions within each risk loci that cumulatively contained 13482 variants (4331 previously published candidate SNPs with PICs and additional 9151 SNPs in high LD in our sequences). Out of the total 13482 variants considered, we were able to map 9372 on the RELATE-inferred local trees such that each tree branch contained one SNP (4838). The rest of the SNPs did not pass QC requirements specific to the RELATE approach. Next, for each inferred tree, we estimated the log-likelihood ratio for the positive selection scenario versus the neutral scenario using the CLUES approach. The overall RELATE and CLUES-based analyses consist of several key steps briefly outlined below.

To test candidate SNPs for signatures of positive selection, we start by building local genealogies for whole-genome sequences from Estonian Biobank using Relate ^48^; We then extract subtrees corresponding to samples of interest. Next, based on the extracted subtrees, we estimate the coalescence rate through time. The coalescence rate through time is used to trace the trajectory of the effective population for the Estonian population. This information is needed for CLUES analysis and to mimic the demographic history for the Estonian population when we simulate sequences and explore the logLR distribution expected under neutral demography. Finally, we run the CLUES inference for the SNPs of interest using the extracted subtrees and coalescence rate through time. Below, we describe each step in the RELATE and CLUES pipeline in more detail.

### Overview of our approach to detect positive selection

Next, we aimed at testing for evidence of natural selection acting on the SNPs selected above. This section provides an overview of our approach, while technical details are given in the corresponding sections below.

We chose CLUES ^28^ as a tool for detecting positive selection as it has several desired properties: 1) it returns estimates of both the strength of selection in the form of the selection coefficient (s) and the support for a non-neutral change in allele frequency at the focal SNP in the form of a log-likelihood ratio (logLR) between the model with the inferred value of s and a neutral model (s = 0); 2) comparison of logLR values between partially linked SNPs can be used for selection signal fine-mapping; 3) it allows the user to specify the time period to focus on and 4) it is suitable for detecting selection on standing variation. This is possible because CLUES estimates the derived allele frequency trajectory over time for a tested SNP using the local tree’s topology and branch length (BL) as input. A local tree represents evolutionary relationships (the order and time of coalescence) between DNA sequences at a given genomic location. As we move along the chromosome, the topologies and BL of the local trees change as we cross points of recombination events that took place during the history of the studied DNA sequences; however, overall, neighbouring trees are highly correlated in their structure. The properties of a local tree are disturbed by positive selection in a predictable manner, resulting in an increased coalescence rate (and hence shorter length) of branches carrying the selected allele within the time interval when the selection was acting. The local trees and the genome-wide average estimate of coalescence rate through the time required by CLUES as input can be obtained by running Relate - a local tree inference tool that relies on whole-genome sequences ^48^.

Thus, our inference consisted of the following steps:

1. Preparing the whole-genome sequences data set and applying Relate ^48^ to it to build local trees.
2. Extracting subtrees corresponding to a subset of samples to a) filter samples for CLUES and b) use a smaller subset for estimating coalescence rate over time which becomes computationally costly with too many samples.
3. Running the coalescence rate / BL estimation procedure on a small subset of the samples but on the whole genome to obtain the neutral expectation of coalescence rate over time.
4. Extracting a local tree corresponding to a given SNP of interest and resampling its BL to take into account the uncertainty in BL estimate. This results in a sample of local trees with the same topology but different BL. As each new tree in the sample is obtained by re-estimating the BL of the previous tree, the earlier trees in the sample are likely to have less accurate BL, and hence it makes sense to remove those as burn-in. Also, neighbouring trees in the sample are likely to have similar BL estimates, and hence it is desired to “thin” the sample of trees to avoid redundancy by keeping every i^th^ tree.
5. Running CLUES on the sample of local trees for a given SNP obtained from the previous step.

Steps 1-3 were run once for the whole genome, while steps 4 and 5 were run independently for each SNP of interest. As described above, we started with 13482 variants. Out of those 9372 passed the applied mask (see below) and were mapped to RELATE-inferred local trees and thus could be used for CLUES. However, as several SNPs can map to the same branch of the same local tree (reflecting perfect linkage), thus providing the same information, we analyzed one SNP per branch (4838 in total) and then assigned the same value to all SNPs residing on the same branch.

Note that the overall approach we use here to run CLUES is the same as in Marnetto et al, 2021.

### Tree building

The local genealogical trees were built by applying Relate version v1.1.4 to 2420 phased whole-genome sequences of the Estonian Biobank participants described in Kals et al., 2019 and Pankratov et al., 2020. We kept only SNP positions with high variant calling certainty in our sequence data using the strict callability mask (GRCh37) from the 1000 Genomes project (The 1000 Genomes Project Consortium). SNP alleles were polarised into ancestral and derived based on the GRCh37_e71 homo sapiens ancestral genome, and SNPs with unknown ancestral stater were removed. To carry out the tree-building procedure, we used the GRCh37 recombination map (The 1000 Genomes Project Consortium), the mutation rate of 1.25×10-8, and the effective population size of 30000 haploids.

### Extracting subtrees

Since CLUES expects a panmictic population, we removed close relatives, ancestry outliers, and clusters of individuals with excessive IBD sharing. For this, we started with the dataset of 2305 samples described in Pankratov et al., 2020, where they removed extreme PCA outliers and close relatives up to 3rd degree. In this dataset, we additionally removed a) outliers in PCA within the Estonian dataset or when projecting Estonian samples on the PC space defined by samples from various European populations; in both cases, we removed samples falling out of the 2.5-97.5 percentile range for the first and second Principal Components; b) top and bottom 2.5% of the samples based on singleton counts per genome and c) samples that had pairwise total IBD sharing of 166.2 cM or more with more than one another sample in the dataset. All the analyses used for these filtering steps are described in Pankratov et al., 2020. After filtering, we randomly subsampled 1800 individuals for CLUES. We further downsampled these 1800 to 100 individuals for coalescence rate through time estimation.

Therefore, the subtrees for the subset of 100 and 1800 samples were extracted separately using the SubTreesForSubpopulation mode in the RelateExtract program.

### Estimating coalescence rate through time

To estimate the coalescence rate through time, we applied Relate’s EstimatePopulationSize.sh module to the random set of 100 samples with the following parameters: mutation rate 1.25×10-8, generation time 28 years, number of iterations 5, tree dropping threshold 0.5, and time bins defined as 10x years ago where x changes from 2 to 7 with an increment of 0.1.

### CLUES analyses

Next, we used Relate’s local trees corresponding to the 1800 individuals as input for CLUES selection analysis. For a target SNP, we extracted the corresponding local tree and resampled its branch length 200 times. We removed the first 100 sampled trees as burn-in and pruned the remaining 100 trees by keeping every 5^th^ tree. The resulting 20 trees were then used for CLUES. We focused on a time period between 0 and 500 generations ago. As CLUES relies on sampling branch length in an MCMC-like manner, it does not return the same result when run on the same SNP twice, and the degree of discordance depends on the uncertainty in branch length, which is tree-specific. To account for that, we ran CLUES twice for each SNP (starting from BL sampling), and if the logLR values differed by more than two units (indicating higher uncertainty), we ran it for the third time. Then we assigned the median values of logLR and s (out of 2 or 3 observations) to each SNP.

### Distribution of LogLR values expected for Estonian population demography

We next performed a simulation study to assess the LogLR distribution expected for the Estonian population history. We first used the coalescence rate through time estimated using Relate as described above to get the effective population size (Ne) trajectory for the Estonian population (Supplementary Table 3 and Supplementary Fig 2a). We then used this Ne trajectory to inform parameters choices to simulate 3600 sequences (equivalent to 1800 diploid individuals) corresponding to chromosome 1 (GRCh37) in length using msprime (Kelleher et al., 2016). We used recombination rates along the chromosome extracted from the GRCh37 recombination map of chromosome 1 (The 1000 Genomes Project Consortium) and a mutation rate of 1.25×10^−8^ per nucleotide per generation. To simulate demographic events at generations 0 to 1000, we used the discrete-time Wright-Fisher model (model=“dtwf) and then switched to the Hudson model (model=“hudson”) after generation 1000. The simulated sequences were then used to perform tree-building (for 1800 diploids) and coalescent rate through time estimation (for a random subset of 100 diploids) as described for the real data. The resulting Relate output was then used to perform CLUES analyses. To select SNPs for the CLUES analyses, we first removed SNPs that matched at least one of the following criteria: a) not passing the strict callability mask, b) not mapping to the inferred trees, and c) had minor derived allele count less than four in the sample (1800 diploids), which resulted in 765217 SNPs, altogether. We further pruned our SNP set by retaining every 76^th^ SNP and used the resulting 10,068 SNPs for our CLUES analyses performed as described for the real data.

## Acknowledgements

This research was supported by the European Union through the European Regional Development Fund (Project No. 2014-2020.4.01.16-0125, Project No. 2014-2020.4.01.16-0030, Project No. 2014-2020.4.01.15-0012, MOBEC008), European Union through Horizon 2020 research, and innovation program under grant no 810645, Estonian Research Council (grants PRG243), Russian Foundation for Basic Research (grant 18-04-00972\18), and Government of the Russian Federation through the ITMO Fellowship and Professorship Program to B.Y.

We thank Leo Speidel and Aaron Stern for their guidance and help with running Relate and CLUES analyses. Data analyses for this study were carried out in the High-Performance Computing Center of the University of Tartu.

## Author Contributions

Conceived and designed the experiments: B.Y. M.Y. Analyzed the data: B.Y., V.P. M.Y., and S.R., Interpreted and Discussed results: V.P., M.Y., B.Y. Implemented simulation study: V.P., M.Z. and S.R. Contributed reagents/materials/analysis tools: M.Z., S.R., V.P. Wrote the paper: B.Y., M.Y., and V.P.

## Consortia

Estonian Biobank Research Team:

Andres Metspalu, Mari Nelis, Lili Milani, Reedik Mägi & Tõnu Esko

## Supplementary Material

**Supplementary Figure 1 Schematic overview of the sequence data and analyses**

**Supplementary Figure 2 Relate-inferred history of effective population size (Ne) for Estonian population and simulated logLR values based on that Ne history**

Panel a. Effective population size (Ne) is plotted against time in human generations. Ne values abbreviated as 5K and 1M should be read as 5 thousand (kilo) and one million (mega) individuals, respectively. Line in dark blue shows Ne trajectory inferred from Estonian population using *Relate*. The dashed line in grey shows Ne parameters adapted and used for simulations. Line in light blue shows Ne trajectory inferred from simulated data using *Relate*.

Panel b shows the distribution of logLR from 10000 simulated datasets, and three vertical lines in light blue show 95%, 99%, and 99.9% percentile points, respectively. We use 95% percentile point 1.59 as our neutrality rejection threshold.

**Supplementary Figure 3 Local tree-based analysis distinguishes novel peaks with sweep-driving SNPs in previously reported selection signals**.

Six graphs (from top to down) represent five layers of annotation for each candidate SNP in the analysed risk loci. The first graph on the top shows logLR values. The horizontal dashed line in orange separates logLR values above the neutrality threshold. Red arrows point to the three logLR peaks. The second graph depicts PICs values for each candidate SNP reported in Farh 2015. PICs score measures the probability of the candidate causal variant to be causal. The third graph from the top shows risk haplotypes composed of the marker SNP along with linked candidate SNPs and their effect alleles. When the SNP allele is derived, it is shown in green and blue when ancestral. Three asterisks (^***^) mark the most likely candidate causal SNPs, which has the highest PICs score. The “o” suffix denotes the SNP with evidence for selection (logLR≥1.59), and “ooo” highlights the most likely sweep-driving mutation. Appended letters s+ and s-indicate positive and negative selection coefficients, respectively. The fourth graph shows the heatmap of pairwise linkage disequilibrium between SNPs, measured in r^2^. The fifth and sixth graphs show gene annotations and regulatory elements from the Ensembl database (version 87, Human genome build GRCh37).

**Supplementary Figure 4 LogLR variation around the adaptive rs3184504*A variant in SH2B3 gene**

Six graphs (from top to down) represent five layers of annotation for each candidate SNP in the analysed risk loci. The first graph on the top shows logLR values. The horizontal dashed line in orange separates logLR values above the neutrality threshold. Red arrows point to the three logLR peaks. The second graph depicts PICs values for each candidate SNP reported in Farh 2015. PICs score measures the probability of the candidate causal variant to be causal. The third graph from the top shows risk haplotypes composed of the marker SNP along with linked candidate SNPs and their effect alleles. When the SNP allele is derived, it is shown in green and blue when ancestral. Three asterisks (^***^) mark the most likely candidate causal SNPs, which has the highest PICs score. The “o” suffix denotes the SNP with evidence for selection (logLR≥1.59), and “ooo” highlights the most likely sweep-driving mutation. Appended letters s+ and s-indicate positive and negative selection coefficients, respectively. The fourth graph shows the heatmap of pairwise linkage disequilibrium between SNPs, measured in r^2^. The fifth and sixth graphs show gene annotations and regulatory elements from the Ensembl database (version 87, Human genome build GRCh37).

**Supplementary Table 1 Relate and CLUES-based results for all the tested 9372 candidate SNPs**

**Supplementary Table 2 Candidate causal SNPs in balancing haplotypes reported in Siewert & Voight, 2017**

**Supplementary Table 3 Effective population size (Ne) parameters inferred for Estonian population**

**Supplementary Data 1** accompanies Figure 3 and contains full annotations for the 204 risk loci with logLR, PICs scores, risk haplotype, Ensembl genes and regulatory elements. Available at https://figshare.com/s/c6b80455488afbd1655a

