## Supplementary figures and images for "Prioritising Autoimmunity Risk Variants for Functional Analyses by Fine-Mapping Mutations Under Natural Selection"

### Supplementary Figure 1

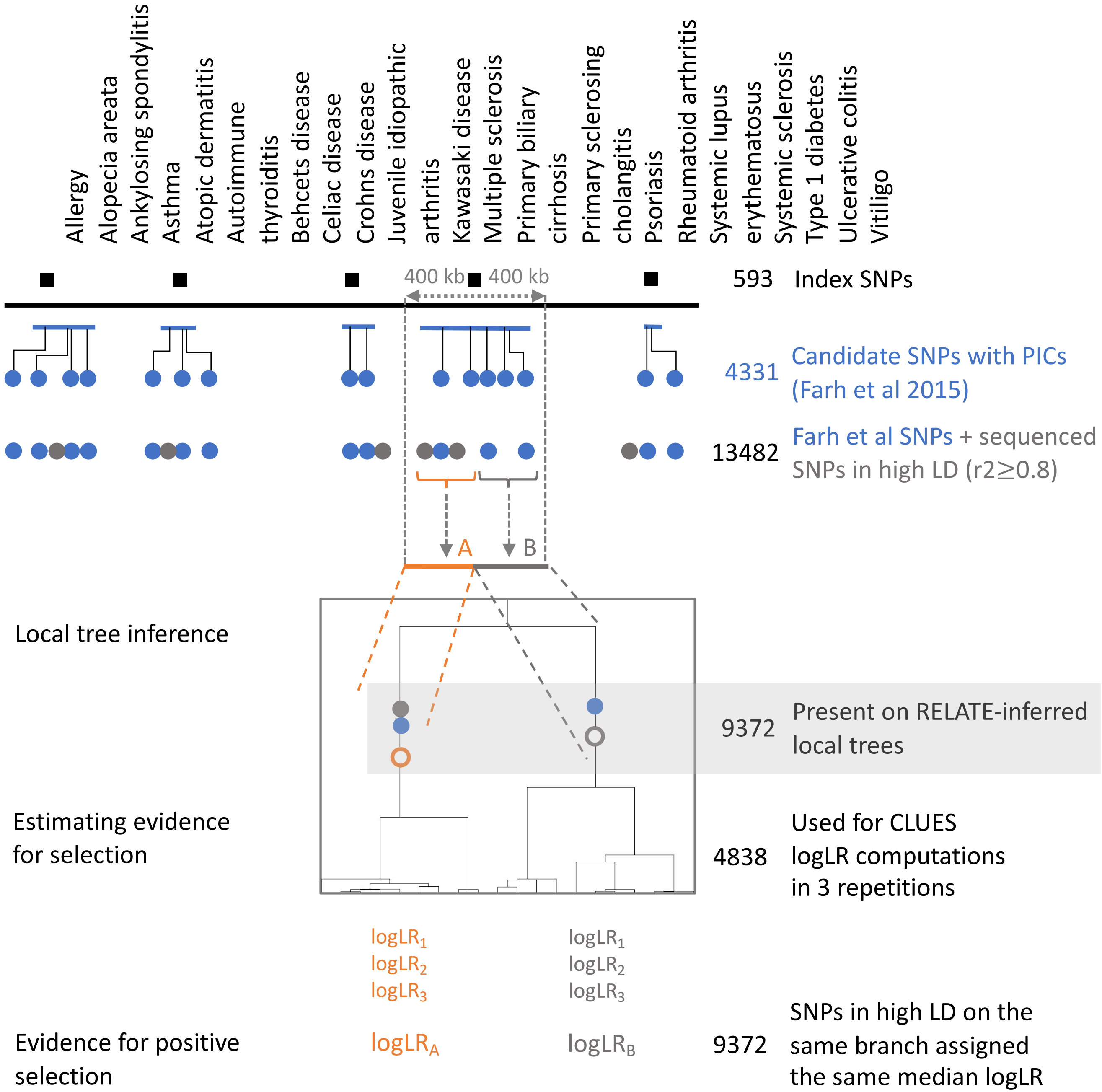

### Supplementary Figure 2

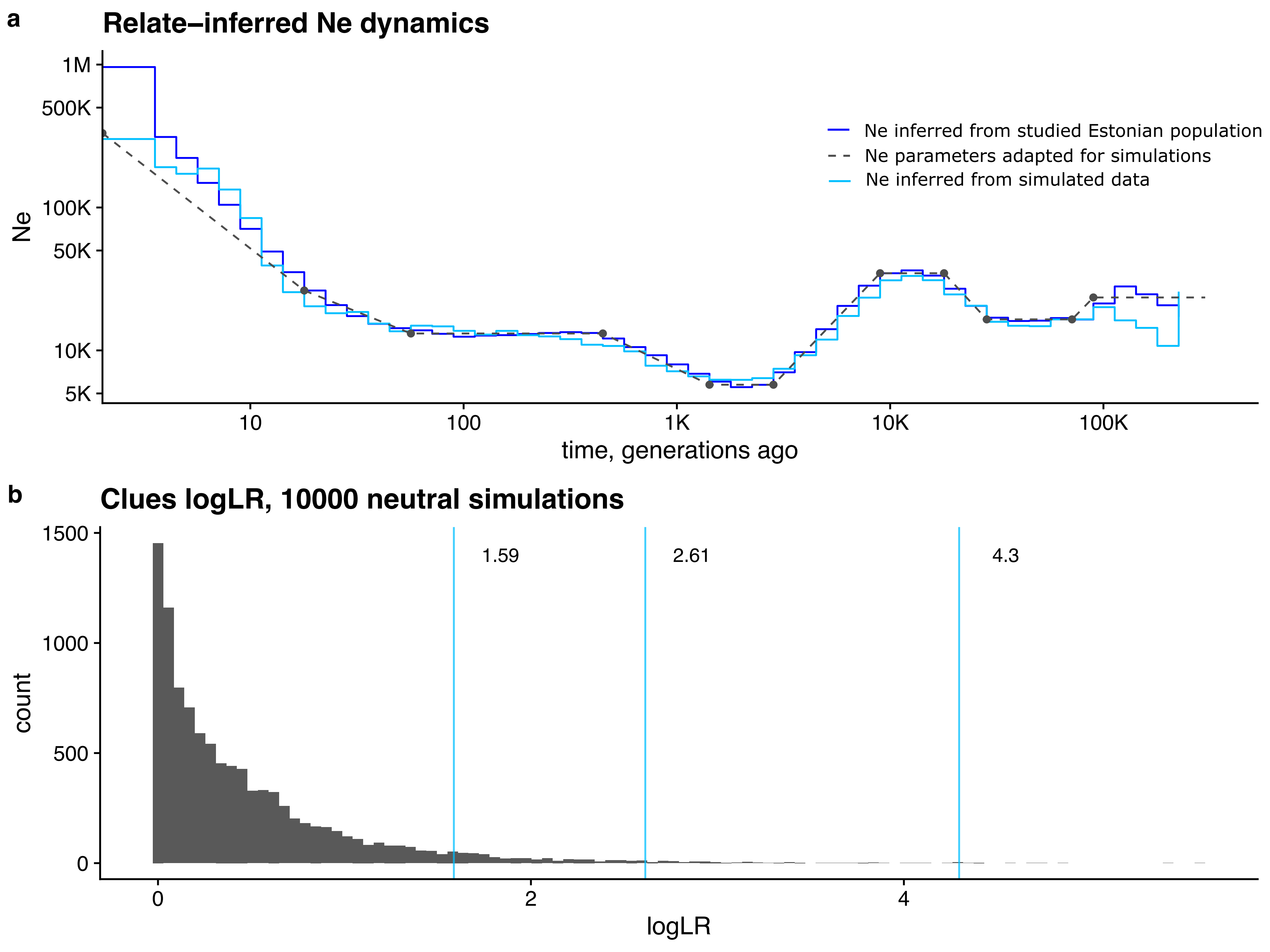

### Supplementary Figure 3

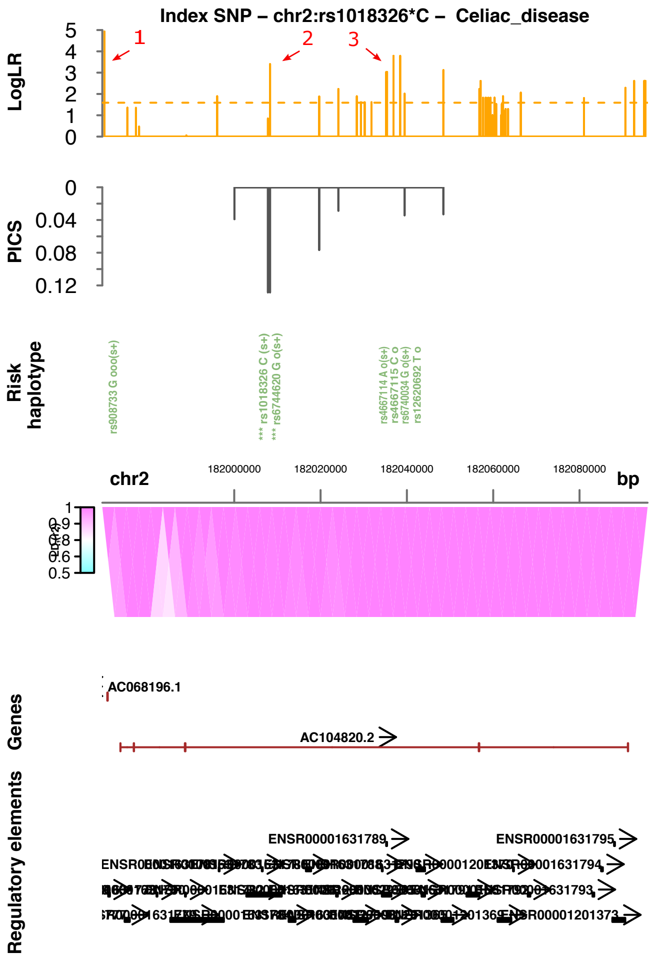

### Supplementary Figure 4

#### Index SNP – chr12:rs3184504*T – Celiac_disease

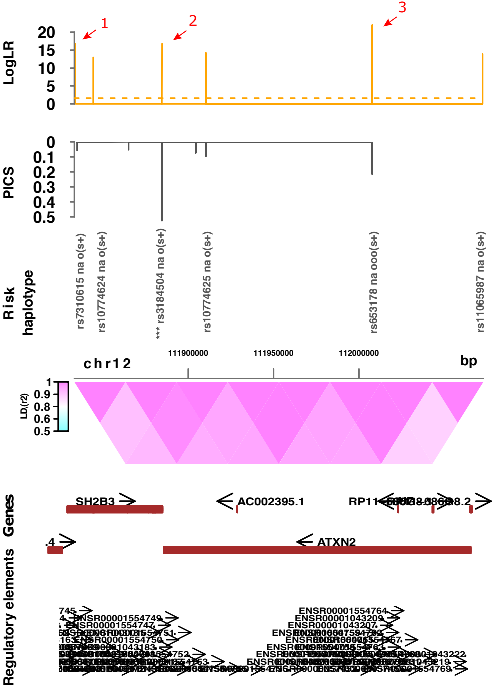
